# NC4touch: An open-source networked touchscreen apparatus for rodent behavioral testing

**DOI:** 10.64898/2026.07.30.741621

**Authors:** Gelareh Modara, Adam W. Lester, Matthew B. Cooke, Isaac Schwein, Vanesse Li, Jocelyn Zhang, Amy Wong, Leticia Cid, Jason Snyder, Manu S. Madhav

## Abstract

In recent years, rodent touchscreen-based paradigms have gained popularity for their flexible task design, automated data collection and improved standardization. These devices minimize experimenter intervention and contribute to more replicable and reliable research. However, the high costs and proprietary hardware / software associated with commercially available systems present a financial barrier for smaller labs or researchers with limited resources. To address this issue, we developed “NC4Touch”, an open-source scalable, modular rodent testing apparatus. It features three independent touchscreens and a feeding port for easy use in a wide array of tests comparable to those available with standard operant chambers, with the added benefit that visual stimuli can be highly customized. The system includes a user-friendly graphical interface that offers real-time control, task customization, video recording, and data management. We demonstrate its effectiveness in a visual discrimination task using both rats and mice. Several devices can be operated in parallel from the same computer and user interface, allowing high-throughput data collection. We provide detailed assembly instructions for the hardware and well-documented and easily-configurable software. We hope that the affordable open-source nature of NC4Touch will expand the scope of behavioral testing, allowing researchers to overcome traditional financial barriers and create a more collaborative community.

**Significance Statement:** Touchscreen-based testing is a powerful tool for assessing cognition in rodent models, however commercial systems remain cost-prohibitive for many researchers. *NC4Touch* is a rodent touchscreen apparatus that provides an open-source, customizable, and affordable alternative that enables high-throughput, cognitive testing in both mice and rats. By lowering financial barriers and promoting hackability, this system enables greater accessibility and collaboration in research.

## 1. Introduction

In recent years, advancements in systems neuroscience have increasingly relied on behavioral assays to investigate learning, memory, and decision-making in animal models. Among these, rodent touchscreen-based paradigms have become quite popular due to their high flexibility in task design, improved standardization across studies, and automated data collection. These factors minimize experimenter intervention, contributing to more replicable and reliable research outcomes^1^. However, commercially available touchscreen apparatuses are costly (>$10,000 USD per chamber), which poses a significant financial barrier for laboratories with limited resources, particularly if their studies call for high-throughput data collection. In addition, commercial apparatuses tend to have closed ecosystems, with non-repairable hardware and minimally configurable software. Given these constraints, there has been growing interest in affordable open-source, do-it-yourself (DIY) touchscreen alternatives that can offer similar functionality as commercialized touchscreens without significant expense. Over the past decade, a few rodent touchscreen chambers have been developed using affordable components, sharing designs and software with the community^2–4^. These DIY platforms can be modified and researchers can program new task variants or tweak parameters as needed. Hardware modifications are also possible; for instance, one can swap in a larger touchscreen or integrate additional sensors (cameras, IR beams, lick detectors). This ‘hackability’ allows scientists to tailor the apparatus to their experimental needs. On the other hand, one must consider that building a DIY apparatus requires at least a moderate level of technical proficiency in design, electronics and programming, as well as access to the necessary fabrication tools. Thus, these are often the domain of multidisciplinary labs with engineering expertise, or labs with sufficient resources to hire dedicated technical personnel.

In addition to fully open-source DIY platforms, custom-built touchscreen systems have also been developed for specific behavioral applications^5^. These systems are typically lower-cost alternatives to commercial touchscreen chambers and further reflect the growing interest in accessible behavioral neuroscience tools. However, many remain specialized toward particular experiments and may not provide comprehensively open-source hardware/software ecosystems for broader community adaptation and scalability. We developed a customizable and cost efficient (∼$550 USD) cognitive touchscreen chamber designed specifically for rodent behavioral experiments. Built using commonly available microcontrollers and hobbyist hardware, this open-source apparatus provides an accessible solution that is comparable in functionality to commercial systems. The design is modular and flexible, allowing users to modify both the hardware and software to accommodate tasks ranging from basic stimulus-response tests to complex visual discrimination paradigms. The networked architecture furthermore enables users to control and monitor multiple chambers from a single computer, enabling high-throughput data collection.

## 2. Materials and Methods

### 2.1. Hardware

A detailed bill of materials as well as assembly guide for the apparatus is available on the Open Science Framework (https://osf.io/t5vfp).

#### 2.1.1. Chamber

The NC4Touch apparatus was designed using Solidworks (v2023, Dassault Systèmes, Vélizy-Villacoublay, France) and was inspired by design principles from previous work^6^, as shown in Fig. 1. The chamber’s interior features a trapezoidal workspace specifically shaped to guide the animal’s attention toward the touchscreens (Fig. 1a) in the front and reward zone in the back (Fig. 1b). The behavioral area measures 260 mm at the touchscreen end, 70 mm wide at the reward end, and 350 mm in depth, with a height of 245 mm and walls ∼3 mm thick. Structural support is provided using 10×10 mm miniature T-slotted framing rails. The walls are constructed from 1/8- inch acetal, while the lid and floor are made from 3/16-inch clear acrylic.

**Figure 1:**
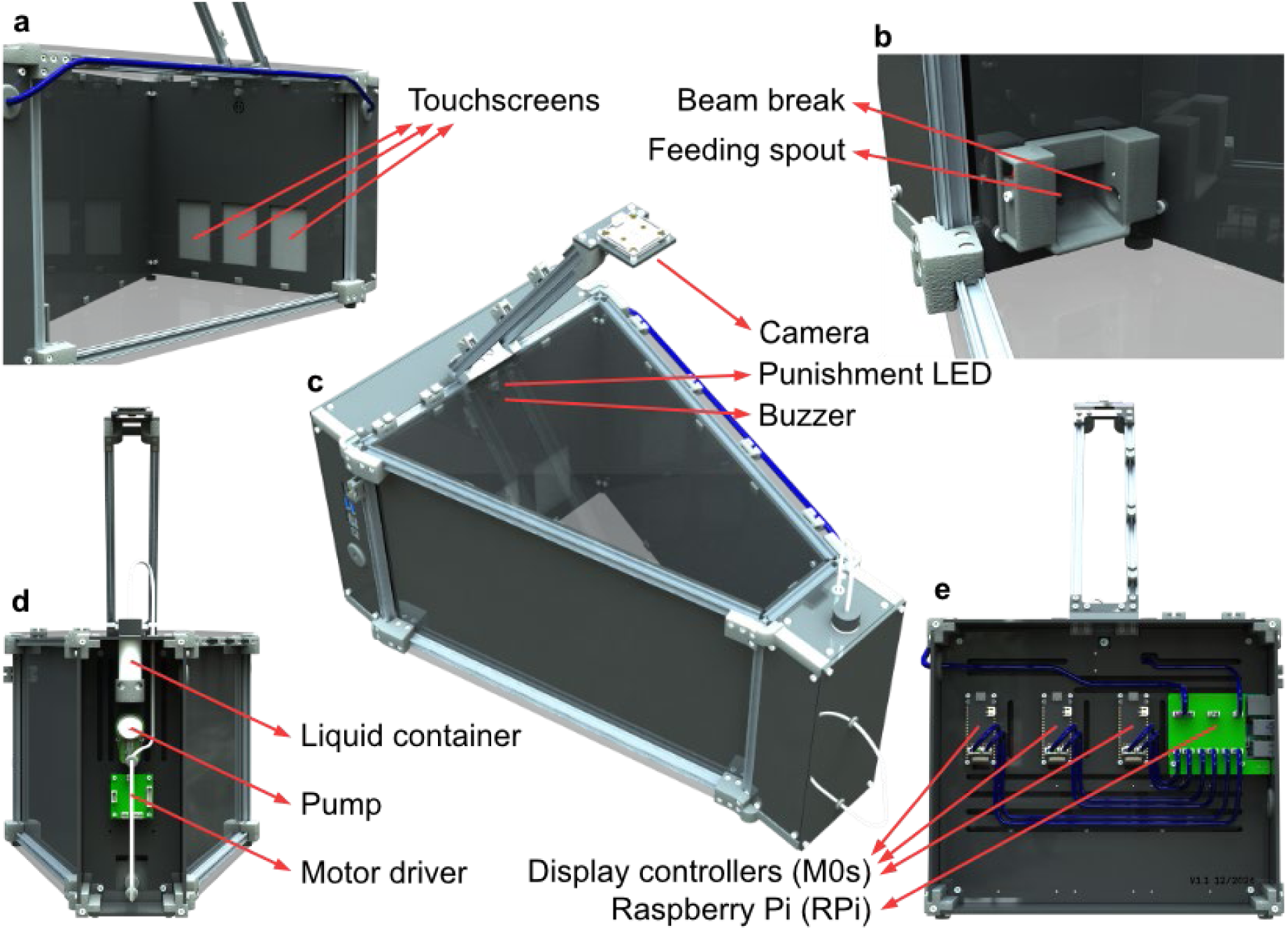
3D Renderings of NC4Touch apparatus. (a) Cutout view of interior front highlighting the touchscreens. (b) Zoomed view of interior back highlighting the reward port. (c) Perspective view showing the whole apparatus. (d) View of exterior back showing the reward system. (e) View of exterior front showing the display and control boards.

#### 2.1.2. Controller

The main controller for the NC4Touch chamber is a Raspberry Pi 4B (Raspberry Pi, Cambridge, UK), hereafter referred to as RPi (Fig. 1e), a single-board computer running the Raspberry Pi OS (Debian Trixie) distribution. This controller allows for both standalone usage of a single chamber, or networked operation of multiple chambers when paired with an additional control computer. The RPi has Ethernet, Universal Serial Bus (USB), Serial Peripheral Interface (SPI) and General Purpose Input / Output (GPIO) interfaces that we use to communicate with peripherals. The RPi boards are also extensible with Hardware Attached on Top (HAT) boards that extend its functionality. For the chamber, we used a GPIO terminal HAT (DFR0918, DFRobot, Shanghai, China) for easy attachment of peripherals, and an optional Power over Ethernet (PoE+) HAT (Raspberry Pi, Cambridge, UK) for combined power and networking over a single ethernet cable.

#### 2.1.3. Camera

The chamber features a raised arm onto which a camera (FIT0729, DFRobot, Shanghai, China) is mounted. This camera has a field of view (76°) that encompasses the entire chamber, and high-resolution video capture (1600 x 1200 px @ 30 fps). The camera is connected to the RPi through USB. To provide consistent illumination for top-down video capture, a 5V chip-on-board LED strip (FIT0876, DFRobot, Shanghai, China) was mounted on the camera stalk and positioned to shine directly onto the chamber. The light is wired to a GPIO port on the RPi so that its brightness can be controlled. For behaviors that require limited illumination, these can be substituted with an infrared camera and illuminator.

#### 2.1.4. Display module

The core interactive components of our touchscreen apparatus are three Display Modules (Fig. 1a,e), each of which consists of a 320×480 pixel capacitive touchscreen (Fermion 3.5”, DFR0669, DFRobot), and a microcontroller (Firebeetle 2 M0, DFR0652, DFRobot) connected to the screen. The screen size (73.44 mm x 48.96 mm) is ideal for small-animal apparatuses, which often contain 2-5 touchscreens next to each other. The screens use capacitive touch, which is more sensitive and requires less force than resistive touch. We further increased the sensitivity of the screens by changing their firmware settings so that it reliably works for both mice and rats. The screens also have an SD card slot that can be used for image storage. The screen can be operated through a single General Display Interface (GDI) ribbon cable that contains wires for commanding the display, touch input and SD card. Using this single cable significantly reduces cable management and user error.

The M0 microcontroller has features that make it ideal for driving the display module. The controller features a built-in slot for the GDI cable. The software on the controller is written using the Arduino (modified C++) programming language, very commonly used in the DIY electronics community. It also has built-in flash storage which can also be used for onboard storage of images. The controllers are connected to the main board using USB. The modularity of the Display Module and extensibility of the USB protocol means that users can add or subtract screens with minimal changes to wiring and software.

#### 2.1.5. Front panel electronics

The front panel enclosure (Fig. 1a,c) houses three Display Modules, a buzzer (DFR0032, DFRobot), and a (typically) reward-indicating RGB LED (620-1121-317F, Dialight, London, UK) positioned at the top, along with the RPi. The modular and enclosed design of the electronics compartment isolates it from the main behavioral chamber, enabling easy removal for cleaning and maintenance. This configuration also prevents potential damage or interference from the animal, such as chewing or tampering with electronic components.

#### 2.1.6. Reward module

The reward module (Fig. 1b,d) is located at the rear of the chamber. The front face of the module carries a horizontal stainless-steel lick spout positioned just above floor level and a 5 mm green panel-mount LED (620-1121-317F, Dialight) centered 227 mm above the spout to cue reward availability. Behind the reward module front panel, separated from the animal by an acrylic divider, the rear compartment houses a peristaltic pump (RP-CIII-10, Takasago Fluidic Systems, Nagoya, Japan) and its dual motor controller (MD1.3, DFRobot). Food-grade silicone tubing (1/16” inner dia.) carries the reward solution from an external reservoir placed at the top of the reward module, through the pump, and directly to the lick spout.

The hardware components of the NC4touch apparatus and their connections are depicted in the schematic in Fig. 2.

**Figure 2:**
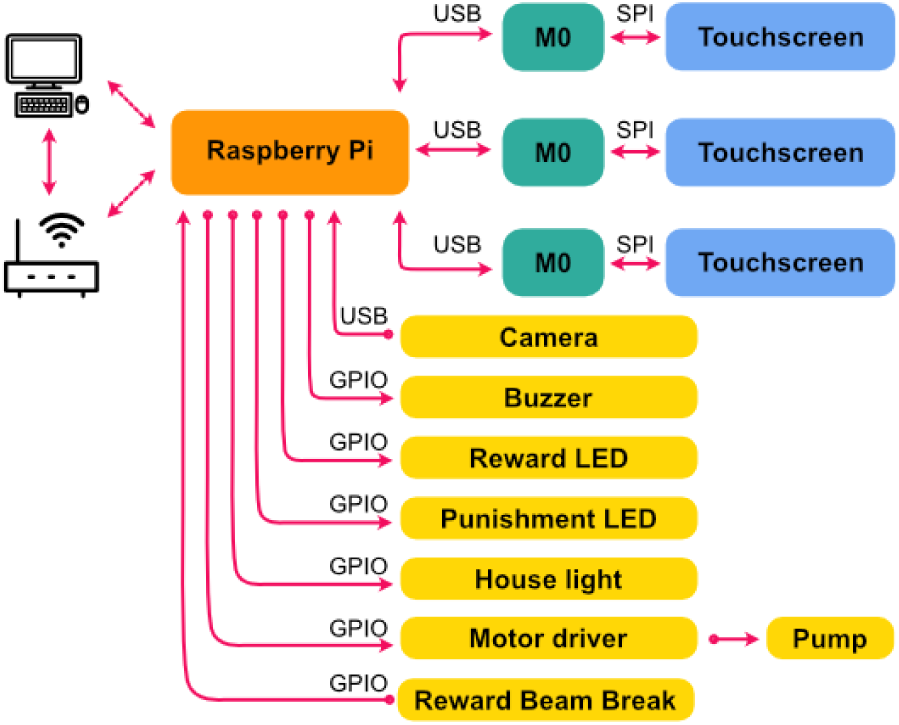
Hardware system schematic. A Raspberry Pi forms the main processing and control unit of the NC4Touch apparatus. Three M0 microcontrollers are connected to the Raspberry Pi through USB link, each of which are in turn connected to a Touch LCD through an SPI link. The Raspberry Pi is also wired up to other peripherals, including the Camera through USB, and other peripherals (Buzzer, Reward LED, Punishment LED, House light, Motor Driver and Beam Break) through GPIO pins.

### 2.2. Software

The NC4Touch software stack is organized as a modular architecture and enables experimental control, hardware management, and behavioral data acquisition. The software architecture consists of five layers: the hardware interface, trainer state machine, session controller, chamber user interface (UI) and network interface. The complete source code for all layers and example behavioral tasks is freely available through the project GitHub repository (https://github.com/NC4Lab/NC4touch).

#### 2.2.1. Hardware interface

Each display module contains an M0 microcontroller running identical firmware that listens for incoming commands from the RPi over USB and controls stimulus presentation on the connected touchscreen. Stimulus images may be stored either on the microcontroller’s onboard flash memory or on SD cards connected to the touchscreens and transferred to the M0 controllers during session initialization. The screens also listen for touches and sends the touch locations back to the RPi as events over USB.

The *μstreamer* package (https://github.com/pikvm/ustreamer) is used to stream camera frames over the local network. Other peripherals (back and front LEDs, house light, reward pump, reward beam break and buzzer) are directly connected to the RPi GPIO pins. These are controlled using the *pigpio* library (https://abyz.me.uk/rpi/pigpio/). We wrote individual Python modules for each of the hardware peripherals so that their operation can be abstracted and simplified when implementing experiment code. A *Chamber* class initializes each of these modules with either default or user-specified values (e.g., GPIO pins used, camera autofocus) that are stored in YAML configuration files on each RPi; this ensures that any configuration updates persist across sessions.

#### 2.2.2. Trainer state machine

Behavioral tasks are implemented in Python as state machines that monitor hardware inputs and control chamber outputs. Each task is encapsulated within a *Trainer* class, responsible for initializing hardware, advancing task logic in response to either time intervals or events and terminating sessions while safely closing data streams. During each cycle, the state machine evaluates hardware conditions and optionally executes a single state transition, such as advancing from stimulus presentation to reward delivery when a correct touch is detected (Fig. 3).

**Figure 3:**
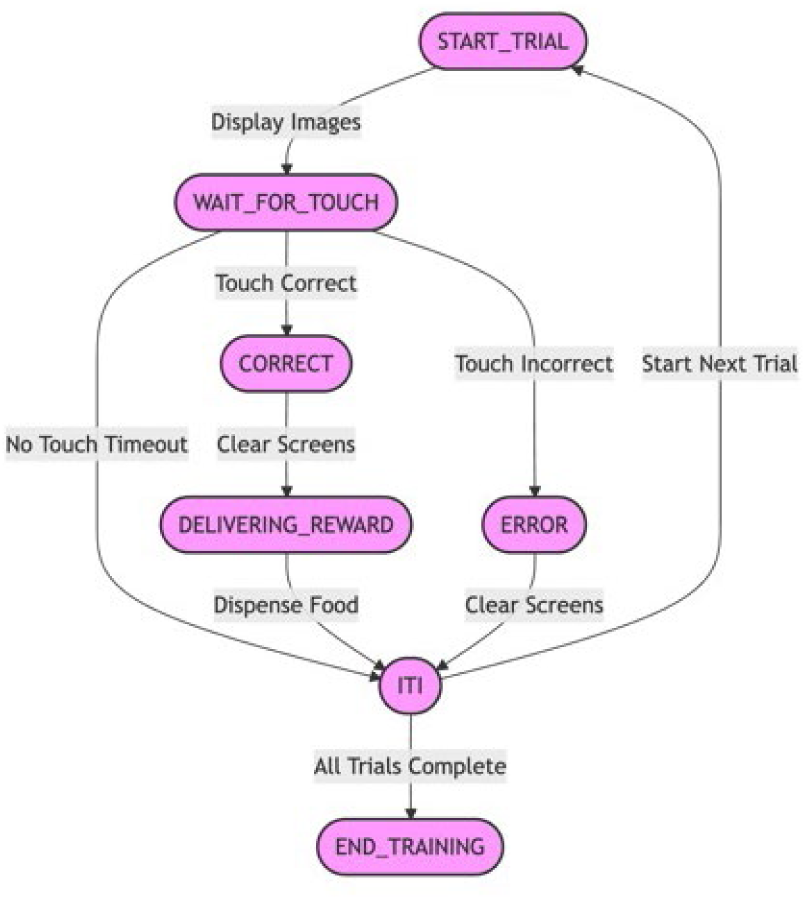
Example Trainer state machine. Each oval represents a state of the algorithm. During each iteration of the training loop, the algorithm either remains in the previous state, or transitions to one of the other states if the conditions indicated on the arrows are met.

#### 2.2.3. Session controller

The session controller functions as the central coordinator between the user interface and the underlying behavioral logic. The *Session* class initializes a *Chamber* class and provides functionality for listing available Trainers, and loading, running, and stopping one *Trainer* at a time. The *Session* also loads the specified experimental configuration, controls experiment timing by executing the training loop at fixed intervals and manages data logging and system shutdown procedures upon task completion. Throughout the experimental session, timestamped trial events are continuously appended to JavaScript Object Notation (JSON) log files to ensure persistent recording of hardware interactions and behavioral data. The complete source code for the M0 firmware, user interfaces, and example behavioral tasks is freely available through the project GitHub repository.

#### 2.2.4. User interface

Experimenters can interact with the system using one of two user interfaces: a terminal-based UI or a browser-based WebUI (Fig. 4a). The selected UI serves as the primary operator dashboard, allowing researchers to configure task parameters, initialize sessions, monitor hardware status in real time, and control session progress. The terminal-based UI provides a simple text interface where the user can monitor and control experiments, whereas the WebUI provides buttons, sliders and other controls to change chamber states and run experiments. The WebUI also enables the user to see the camera feed from the chamber.

**Figure 4:**
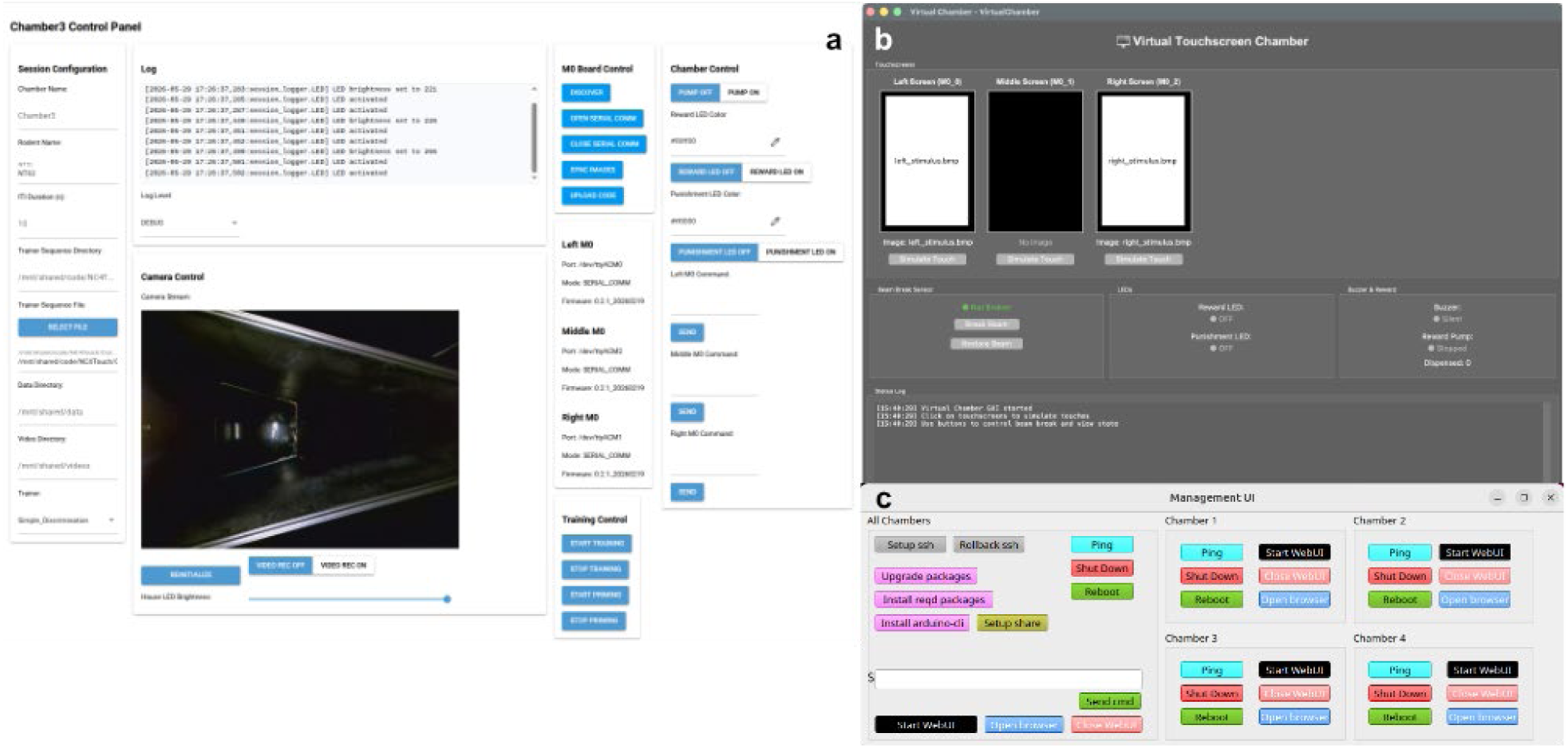
User interfaces. **(a)** The WebUI can be viewed in a browser window, either directly on the Raspberry Pi of each chamber, or on any computer on the local network that the chamber is connected to. **(b)** The Virtual Chamber UI can be used to develop and test training protocols without the chamber hardware. **(c)** The Management UI can setup and control multiple chambers on the local network and allows users to launch their respective WebUIs.

#### 2.2.5. Virtual Chamber

To complement the physical hardware, we developed a virtual touchscreen chamber that closely replicates the behavior of the physical NC4Touch apparatus. The virtual environment was designed to facilitate the development, debugging, and rapid prototyping of behavioral training protocols without requiring access to physical hardware. By simulating the same hardware interface as the physical chamber, the virtualization layer allows existing trainer state machines and behavioral protocols to execute in either physical or simulated environments. Researchers can transition between physical and virtual execution modes through configuration parameters (e.g., a *virtual_mode* flag) without modifying task code.

The virtual implementation consists of simulated hardware modules representing the major components of the NC4Touch system, including touchscreen controllers, infrared beam-break sensors, LEDs, auditory buzzers, and reward delivery systems. Each module maintains and updates its internal state during task execution while generating logged events structurally identical to those produced during *in vivo* experiments. For example, the virtual touchscreen module simulates stimulus presentation by loading bitmap images from local directories and capturing both user-generated and programmatically simulated touch events. Similarly, the virtual reward and beam break modules emulate rodent interactions with the reward receptacle and reinforcer delivery.

The virtual system also includes an interactive graphical interface that visualizes chamber activity in real time (Fig. 4b). Investigators can manually simulate rodent behaviors, including touchscreen responses and reward-port entries, while monitoring hardware states, LED activity, and timestamped system logs. In addition to interactive testing, the framework supports automated unit testing and software validation workflows, enabling rapid verification of behavioral paradigms prior to deployment in animal experiments.

This virtual testing framework allows behavioral tasks to be validated for logical accuracy, state-transition integrity, and hardware synchronization before execution on live hardware. By reducing debugging time and improving software reliability, the virtual chamber substantially accelerates behavioral protocol development and experimental deployment.

### 2.3. Single-chamber Configuration

While operating a single chamber, the Raspberry Pi can be used directly as the experiment computer (Fig. 4A). The experimenter would connect peripherals (monitor, keyboard, mouse) to the Raspberry Pi and operate one of the UIs to run an experimental session. (Fig. 5a).

**Figure 5:**
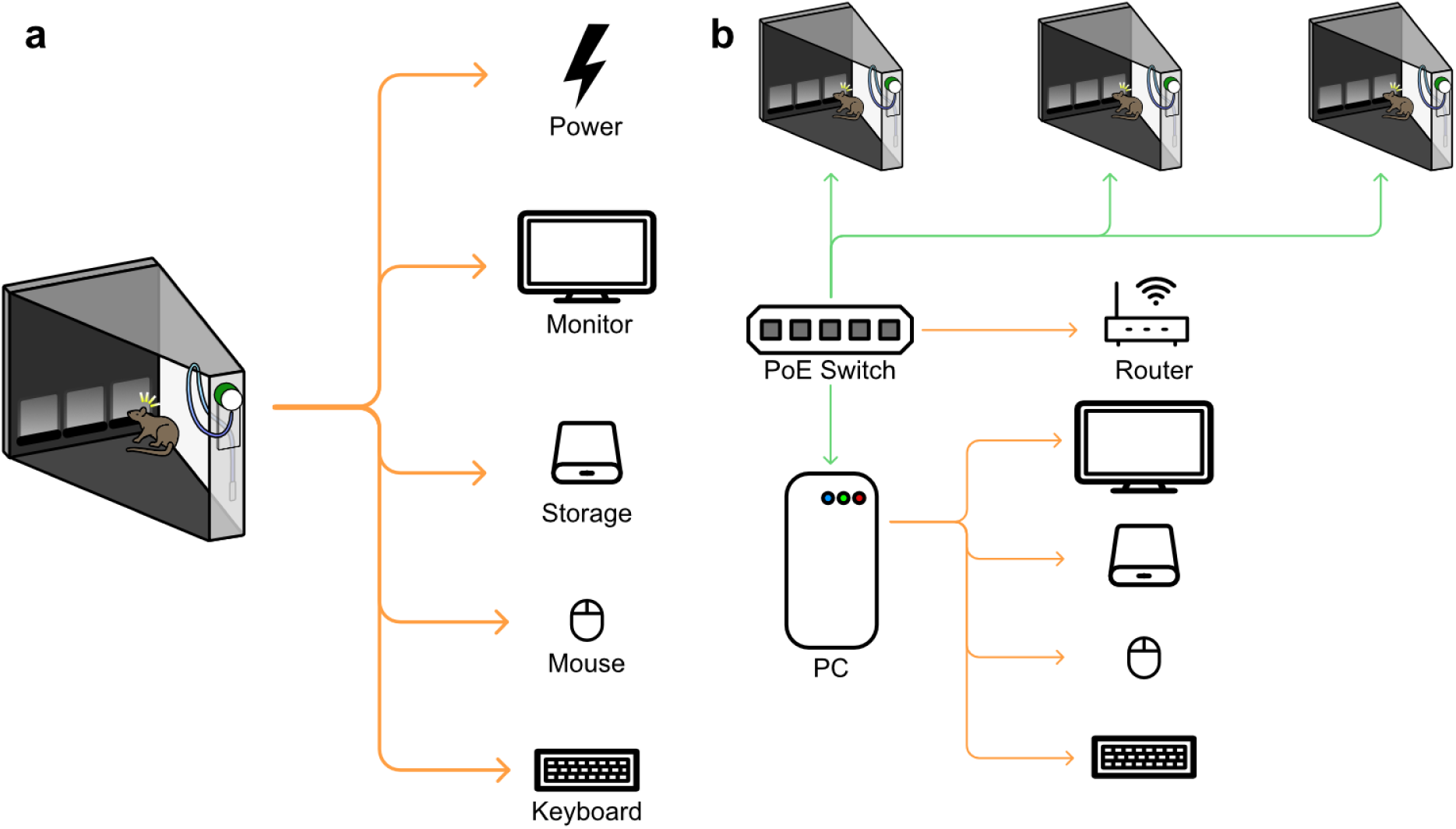
Single- vs. Multi-chamber configurations. **(a)** Single chamber configuration. The RPi on the chamber is directly operated as the control computer – it is wired up to power, input, output and storage peripherals, and any of the three UIs can be used to run an experimental session. **(b)** Multi-chamber configuration. The RPi on each chamber has a PoE+ HAT, and is connected to a PoE switch that provides both power and networking to the chamber. A router connected to the switch manages the network, and the chambers can be controlled using a PC connected to the switch that is part of the same network. Both the browser-based and terminal-based UIs can be used from the PC to run experimental sessions on any of the connected chambers.

#### 2.3.1. Setup

We provide a single script that can be run from a default installation of Raspberry Pi OS (Debian Trixie) that would download and install all required software and dependencies to run the single-chamber configuration.

### 2.4. Multi-chamber Configuration

Multiple NC4Touch chambers can be operated simultaneously. This is accomplished by connecting each chamber to a network switch via an Ethernet (Cat5) cable. (Fig. 5b) Using a Power-over-Ethernet (PoE) add-on board on each chamber along with a PoE-capable switch, this single cable can deliver both power and data to the chamber. A control computer is also connected to the switch, and the experimenter can launch the Management UI (Fig. 4c) on the control computer that can configure the chambers and launch WebUIs (Fig. 4a) to control each chamber through a separate browser window.

A single storage drive on the control computer can be optionally configured to be accessed by all the chambers over the local network. There are several benefits to this: (1) The storage drive can be used to store the code, ensuring that all chambers are running the same code version. (2) The storage drive can store the firmware for the display controllers, and the UI provides an interface to write the firmware to all display modules. (3) The storage drive can hold image files, and the UI provides an interface to transfer these images to the local storage of all connected display modules. (4) The storage drive can be used by each chamber to store experimental data, preventing the need to manually transfer data after experimental sessions.

#### 2.4.1. Setup and Network UI

Once the network setup is wired as shown in Fig. 5b, the user will assign a static IP to each connected chamber. The procedure for how to do this will depend on the network router used. Once this is accomplished, each chamber will have the same IP address every time it is connected to the network.

We recommend that the control computer runs Ubuntu Linux v.24.04. We provide scripts to download and install the required software and dependencies for the control computer, from a fresh operating system installation. Our Management UI (Fig. 4c) then allows users to setup remote access to each connected chamber, install all required software remotely to each of the chambers, run the WebUI server on each of the chambers, and open browser windows to view each of the UIs (Fig. 4a).

### 2.5. Sound attenuation

Scaling the NC4Touch system to a multi-chamber configuration capable of running multiple studies in parallel introduces environmental confounds such as sound interference that may affect behavioral measurements. Notably, leakage of behaviorally-relevant sounds such as the punishment buzzer or reward pump may interfere with rodent task learning. While sound-attenuating chambers are commercially available, often featuring soundproof insulation and ventilation fans (e.g. SmartChamber, Metris b.v., Netherlands), they are typically expensive and harder to access for labs with fewer resources, motivating our development of a low-cost, open-source alternative.

We constructed sound-attenuated boxes (Fig. 6) from medium density fiberboard (MDF) and lined with sound-absorbing polyurethane foam sheets on the top, sides, and door of the enclosure (0.5” thickness, 5692T61, McMaster-Carr, Chicago, USA). The sheets featured a smooth, easy to clean facing and adhesive back which allowed ease of installation. Polyurethane foams facilitate airborne sound absorption by converting sound energy to heat due to friction in the polyurethane cell framework ^7^.

**Figure 6:**
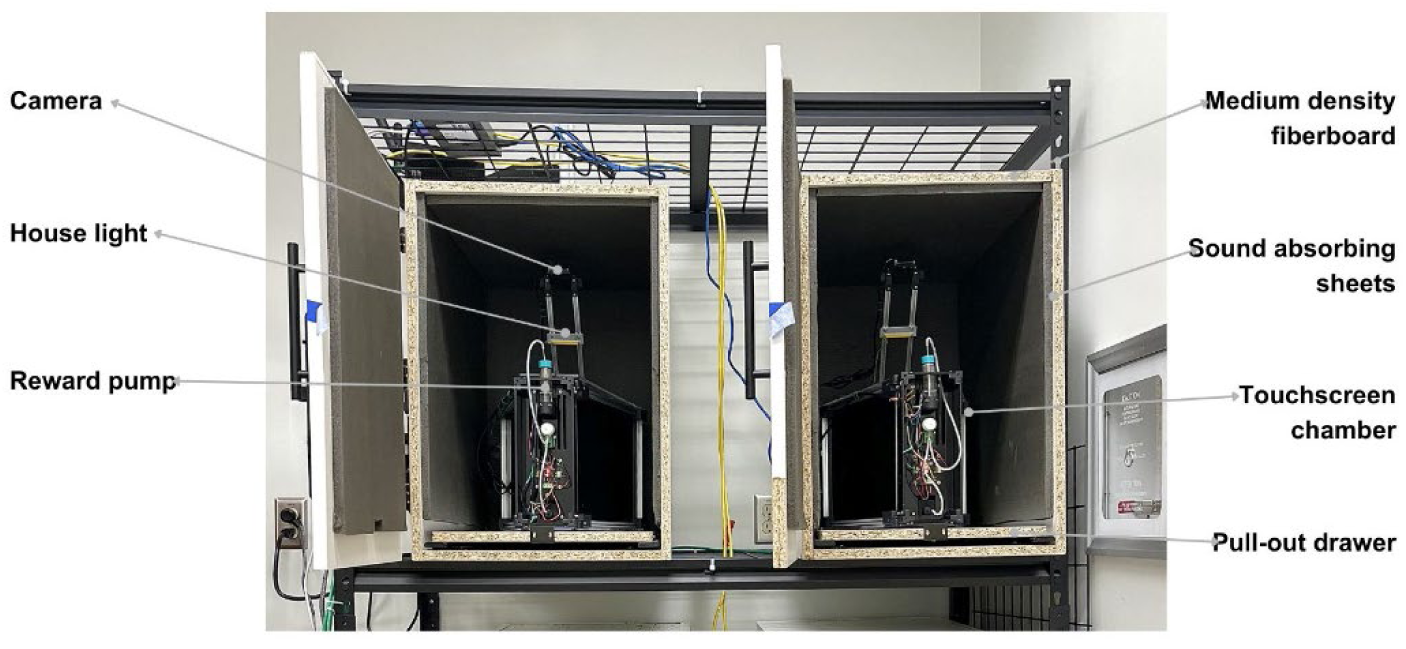
Sound-attenuation chambers. We built MDF boxes lined with sound-absorbing foam and sized them to fit the chamber dimensions. The boxes featured a door for easy access of the chambers from the front, and a pull-out drawer for access of chambers from the top for placing and retrieving rodents. Chambers were inserted with the screen end inside, so that they could be wired from behind and food could be refilled from front.4 boxes were placed in a two-shelf rack, in a 2x2 configuration.

To ensure adequate ventilation and maintain stable temperature and airflow within the chamber, a 5V 90mm fan with a flow rate of 53.38 cfm was installed at the back of the enclosure, oriented to draw air out of the enclosure. The fan was selected for sufficient airflow capacity, providing approximately 582 air exchanges per hour within the approximately 5.50 cubic foot enclosure, and for its ability to be powered directly from the RPi of each chamber. The fan was powered continuously during experiments to prevent heat buildup around the electronics and maintain airflow over the experimental area.

#### 2.5.1. Sound Calibration

The acoustic environment was recorded using an ultrasound microphone (M500-384, Pettersson Elektronik, Uppsala, Sweden). This device converts sound pressure into analog voltage fluctuations, which are digitized internally at 16-bit resolution and 384 kHz before being streamed to the computer. Because the resulting audio samples are expressed in arbitrary digital units, a calibration procedure was required to map raw sample amplitudes to absolute sound pressure level (SPL) values. A portable 1 kHz sound-level calibrator (Model 8930, AZ Instrument Corp, Taichung, Taiwan) was used to record reference tones at 94, 104 and 114dB for offline calibration based on an existing protocol^8^. The root-mean-square amplitudes of these labeled segments were then compared with the known SPLs to derive a calibration constant that permitted conversion of future raw audio traces into calibrated dB SPL.

#### 2.5.2. Sound Attenuation Testing

The main goal of sound attenuation testing was to determine whether the sounds generated by one NC4Touch chamber in normal operation (Chamber A) would be transmitted to an adjacent chamber (Chamber B). A “Sound Test” script cycled through standard NC4Touch events for 60 seconds each while the ultrasonic microphone recorded continuously for five trials. The events were activated and deactivated in the following order: baseline, house light, reward LED, punishment LED, buzzer, touchscreen image display and reward pump.

##### Internal Sound Condition

To quantify the sound produced by a single chamber during normal operation, an ultrasound microphone was positioned inside Chamber A facing the reward pump while Chamber A executed the “Sound Test” procedure. The data files saved for each trial provided a log of each event, including precise information about event names and timestamps.

##### Box-open and Box-closed Conditions

We compared sound transmission between box-closed and box-open conditions to approximate sound levels with or without the barriers of the sound-attenuated box. For both box-open and box-closed conditions, the ultrasound microphone was placed within Chamber B while Chamber A executed the “Sound Test” script. Peak sound pressure levels and frequency spectra recorded in Chamber B during Chamber A task events were compared between box-open and box-closed conditions to quantify inter-chamber sound attenuation. All recordings were analyzed across both audible and ultrasonic frequency bands from 200 Hz to 90,000 Hz.

##### Data analysis

MATLAB version R2025b (Mathworks, Natick MA, U.S.A.) was used to visualize the effects of sound attenuation. To synchronize NC4Touch events with the audio recording, event timestamps were extracted from the chamber data file and recorded audio files. The mean and standard deviation of sound pressures across five trials were calculated for each event and condition to produce a comparison plot. Sound attenuation was quantified as the difference in mean dB SPL between box-open and box-closed conditions for each event. To assess whether these differences were statistically significant, a paired samples t-test was conducted for each event. Cohen’s d was additionally calculated as a measure of effect size.

### 2.6. Behavioral Training

We trained both rats and mice using the NC4Touch apparatus to perform a visual discrimination task. Rat training was done using a single chamber using an earlier version of the software, while mouse training was done using a networked set of four chambers. The protocol and visual cues used also differ slightly. Due to these factors, our goal is not to compare performance between the two species, but simply to demonstrate the utility of our chambers to train either species. All procedures were approved by the Animal Care Committee at the University of British Columbia and conducted in accordance with guidelines set by the Canadian Council on Animal Care.

#### 2.6.1. Rat Training Protocol

Two male and one female Long Evans rats were obtained from Charles River Laboratory. They were single-housed before experiments began and provided *ad libitum* water access. They were then food-restricted, maintaining their body weight at approximately 90% of their *ad libitum* weight.

The training protocol was adapted from the protocol for Pairwise Visual Discrimination (SOP mPVD-v3)^9^ originally created by the Prado Lab and later modified by BrainsCAN Rodent Cognition Core. This protocol was also inspired by previously established touchscreen methodologies for rodent cognitive testing^6,10^, and consists of 8 training stages as summarized in Table 1 and described below. Each training stage was implemented as a training script and deployed on a single NC4Touch chamber.

**Table 1:**
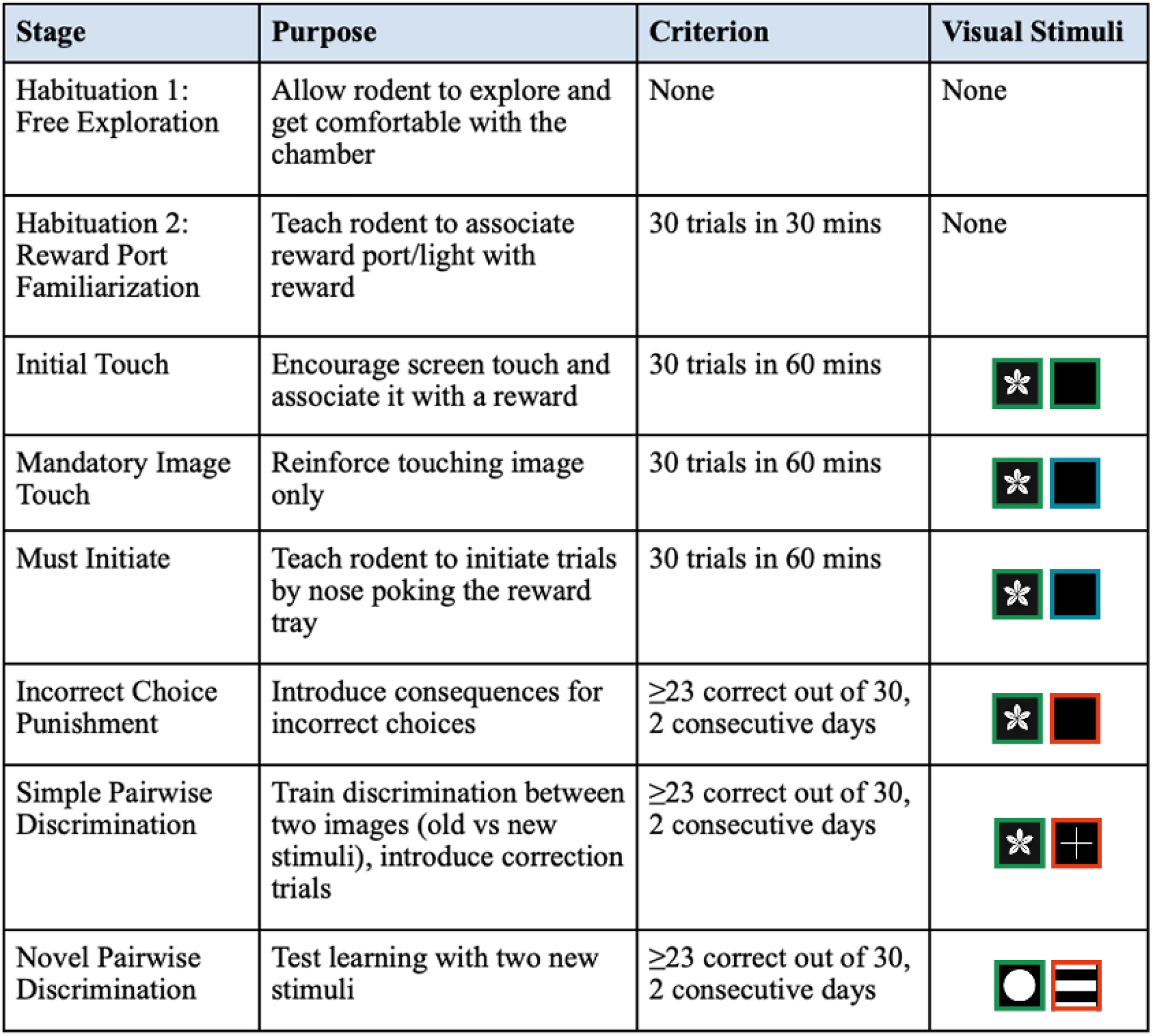
Rat training protocol. Training rats on the pairwise discrimination task required 8 sequential training stages from the naïve animal. The training stages, purposes, graduation criteria and visual stimuli used are detailed in this table.

| Stage | Purpose | Criterion | Visual Stimuli |
| --- | --- | --- | --- |
| Habituation 1:<br>Free Exploration | Allow rodent to explore and get comfortable with the chamber | None | None |
| Habituation 2:<br>Reward Port Familiarization | Teach rodent to associate reward port/light with reward | 30 trials in 30 mins | None |
| Initial Touch | Encourage screen touch and associate it with a reward | 30 trials in 60 mins |  |
| Mandatory Image Touch | Reinforce touching image only | 30 trials in 60 mins |  |
| Must Initiate | Teach rodent to initiate trials by nose poking the reward tray | 30 trials in 60 mins |  |
| Incorrect Choice Punishment | Introduce consequences for incorrect choices | $\geq 23$ correct out of 30, 2 consecutive days | |
| Simple Pairwise Discrimination | Train discrimination between two images (old vs new stimuli), introduce correction trials | $\geq 23$ correct out of 30, 2 consecutive days | |
| Novel Pairwise Discrimination | Test learning with two new stimuli | $\geq 23$ correct out of 30, 2 consecutive days | |

##### Habituation 1, Free Exploration

The rat is placed in the chamber for three consecutive days to allow them to explore and become comfortable in the apparatus, beginning with a 10-minute session on day 1, 15 minutes on day 2, and 20 minutes on. Before each session, liquid reward (∼250 μl) is placed in the reward tray to encourage exploration. This training stage allows rats to explore and become comfortable in the touchscreen chamber.

##### Habituation 2, Reward Port Familiarization

The food tray is primed with a liquid reward (4s, ∼100 μL), and the food tray light is turned on. After the rat enters the reward port, the tray light is turned off. Once the rodent consumes the reward and leaves the port, a 10 s inter-trial interval (ITI) begins. If the rodent remains in the port at the end of the ITI, 1s of delay is added for each additional second the rodent stays in the port. After the ITI, the tray light is turned on again and a reward (3s, ∼75 μl) is delivered. The procedure repeats 30 times. This training stage teaches rats to associate the reward port and its light with receiving a liquid reward. Criterion is reached after completion of 30 trials within 30 minutes.

##### Initial Touch

The food tray is primed with liquid reward (4s, ∼100 μl). One image (e.g., a flower) appears randomly on one of the screens, while the opposite side remains blank. A touch on the image triggers a large reward (4s, ∼100 μl), and a touch on the blank side results in a smaller reward (3s, ∼75 μl), with the reward light turning on in both cases. After the rodent pokes the reward port, the reward light is turned off. Once the rat consumes the reward and leaves the port, a 10 s ITI begins. The position of the images is chosen pseudorandomly such that the stimulus **is not** displayed in the same position more than 3 times in a row. This stage encourages rats to touch the touchscreens and associate screen contact with a reward. Criterion is reached after completion of 30 trials within 60 minutes.

##### Mandatory Image Touch

This is similar to the Initial Touch stage, except that a touch on the blank side results in no response. Only a touch on the image triggers reward delivery with the reward light turning on. The stimuli stay on the screen until the image is touched. This training stage reinforces that only touching the image results in a reward, and that blank touches do not yield any reward. Criterion is reached after completion of 30 trials within 60 minutes.

##### Must Initiate

This is similar to the Mandatory Image Touch stage, except that after the ITI finishes, the food tray light turns on, requiring the rodent to nose poke the reward tray and then exit it before an image is displayed. This training stage teaches rodents to nose poke the reward port before each trial begins, establishing a clear trial initiation signal. Criterion is reached after completion of 30 trials within 60 minutes.

##### Incorrect Choice Punishment

This is similar to the Must Initiate stage, except that touches on the blank side trigger a 5-second punishment light timeout and a 1-second buzzer sound, with no reward delivered. This stage increases discrimination accuracy by introducing a punishment for selecting the blank side. Criterion is reached on 23/30 correct trials (77%) within 60 minutes.

##### Simple Pairwise Discrimination

This is similar to the Incorrect Choice Punishment stage, except that instead of a blank screen, an alternative stimulus (e.g., a cross) is presented. Correction trials are also introduced here: if the incorrect stimulus is chosen, then following punishment and the ITI, the next round will repeat the same trial. The maximum number of correction trials were adjusted based on the researcher’s preference. Criterion is reached on 23/30 correct trials (77%) within 60 minutes.

##### Novel Pairwise Discrimination

This is similar to the Simple Pairwise Discrimination stage, except that two novel stimuli (e.g., a circle and three stripes) are used. This training stage teaches rodents to generalize the task such that one of each pair of stimuli is ‘correct’ and the other is ‘incorrect’. Criterion is reached on 23/30 correct trials (77%) within 60 minutes.

Training progression was monitored through predefined pre-training and fixed criterion per stage. Performance was quantified by the percentage of correct responses per session and the proportion of correction trials. Animals were required to complete a minimum of 30 trials within a 60-minute session, and to reach ≥77% correct choices over two consecutive days in the last three stages to meet performance criteria ^11^.

#### 2.6.2. Mouse Training Protocol

We tested the abovementioned protocol on a cohort of mice – however, they were under a novel water restriction regime that had to be constantly adjusted, and we found their motivation to perform the touchscreen task to be lacking. In our first cohort, only 2/6 mice reached criteria (data not shown). We subsequently reverted to the same food restriction and reward as the rats, as well as made minor modifications to the protocol to optimize training efficiency and task performance.

C57BL;B129S6 mice were purchased from the Jackson Laboratory and initially housed at the UBC Centre for Disease Modelling, before being transferred to the UBC Preclinical Discovery Centre at around 15 weeks of age. All mice were 25 weeks old at the start of the experiment (4 males, 4 females). Animals were dual-housed after transferring to Preclinical Discovery Centre and remain dual-housed throughout testing. These animals were previously tested on a Novel Object Recognition test from week 20-25 of age. The mice were housed on a 12h light/dark cycle with lights on at 7am. Mice always had *ad libitum* access to water. Prior to training, they were food restricted to maintain their body weight at approximately 85% of *ad libitum* weight. Mice were weighed daily and closely monitored in accordance with the Charles River growth chart. Their feeding amount was slightly adjusted according to their rate of weight loss. If an animal fell below 80% of their initial rate, they will return to *ad libitum* feeding until their weight stabilized. Once all the mice’s weight stabilized, weighing took place twice a week instead of daily.

The protocol consists of 8 distinct stages (Summary in Table 2) to gradually encourage rodent interaction with the touchscreens and shape the visual discrimination behaviour. While the stages were very similar to the rat protocol above, we describe them again below for clarity. From stage 3, sessions consisted of 30 trials or a maximum duration of 60 minutes. A 10s inter-trial internal (ITI) followed each trial, during which the house light was dimmed; house light returned to normal brightness at the start of the next trial.

**Table 2:**
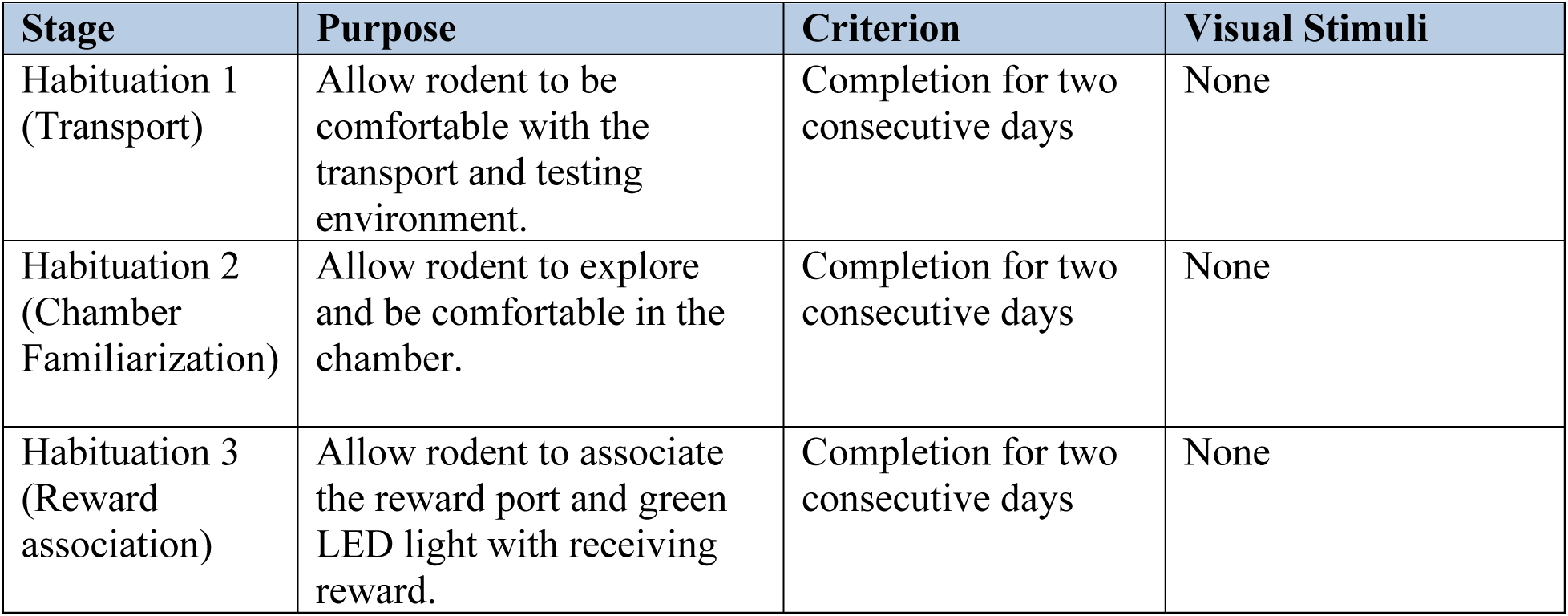
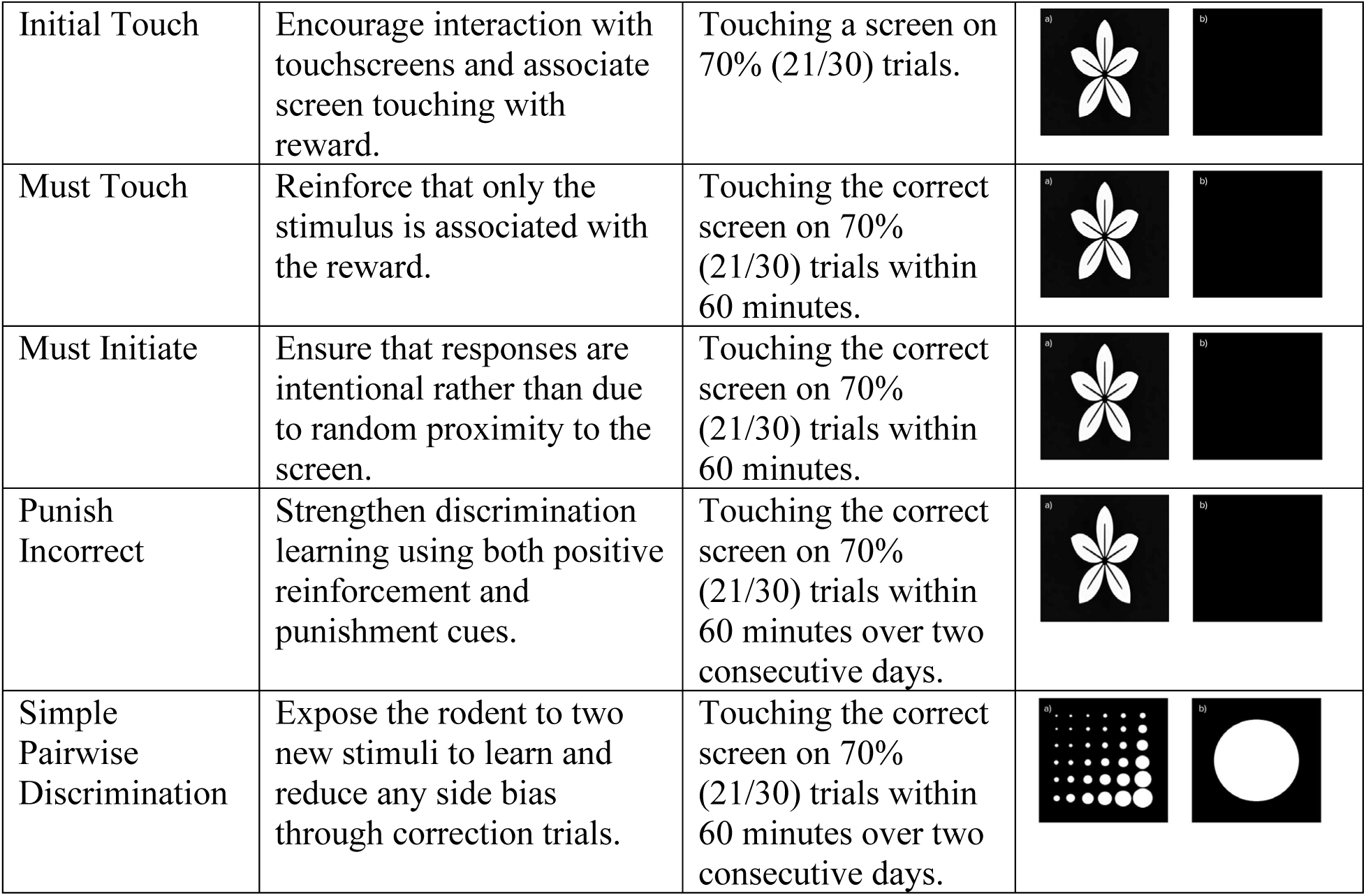
Mouse training protocol. Training mice on the pairwise discrimination task required 8 sequential training stages from the naïve animal. The training stages, purposes, graduation criteria and visual stimuli used are detailed in this table.

| Stage | Purpose | Criterion | Visual Stimuli |
| --- | --- | --- | --- |
| Habituation 1 (Transport) | Allow rodent to be comfortable with the transport and testing environment. | Completion for two consecutive days | None |
| Habituation 2 (Chamber Familiarization) | Allow rodent to explore and be comfortable in the chamber. | Completion for two consecutive days | None |
| Habituation 3 (Reward association) | Allow rodent to associate the reward port and green LED light with receiving reward. | Completion for two consecutive days | None |
| Initial Touch | Encourage interaction with touchscreens and associate screen touching with reward. | Touching a screen on 70% (21/30) trials. |  |
| Must Touch | Reinforce that only the stimulus is associated with the reward. | Touching the correct screen on 70% (21/30) trials within 60 minutes. |  |
| Must Initiate | Ensure that responses are intentional rather than due to random proximity to the screen. | Touching the correct screen on 70% (21/30) trials within 60 minutes. |  |
| Punish Incorrect | Strengthen discrimination learning using both positive reinforcement and punishment cues. | Touching the correct screen on 70% (21/30) trials within 60 minutes over two consecutive days. |  |
| Simple Pairwise Discrimination | Expose the rodent to two new stimuli to learn and reduce any side bias through correction trials. | Touching the correct screen on 70% (21/30) trials within 60 minutes over two consecutive days. |  |

##### Habituation 1, Transport

Mice are transported from the housing facility to the testing room on two consecutive days. They remained in their home cages. This stage allows mice to be comfortable with the transport and testing environment.

##### Habituation 2, Chamber familiarization

Mice are placed in the chambers for 10 min per day over two consecutive days. No stimuli or rewards are presented, and the chambers remain inactive. This allows the mouse to explore and be comfortable in the chamber.

##### Habituation 3, Reward association

At the start of each trial, a milkshake reward (2s., ∼50 μL) is dispensed into the reward port, and the green LED and house light illuminated. Mice are required to collect the reward within 10s; otherwise, the trial automatically enters ITI. After reward collection, another 10 s ITI begins. If the mouse pokes the reward port during ITI, an extra second is added to the ITI duration, up to a maximum of 20 s. 30 trials/day were ran for two days.

##### Initial Touch

A visual stimulus (flower) appears on one of the screens at random, while the opposite screen remains blank. On correct response, a big reward is delivered (3 s, ∼75 μl), followed by a 10 s ITI after reward collection. On incorrect response, a small reward is delivered (1.5 s, ∼37.5 μl) followed by a 10s ITI. If no response is made within 2 minutes, the trial ends and enters ITI. This stage encourages rodents to interact with the touchscreens and associate screen touching with a reward. Criterion is reached when the mouse touches the screen 70% of the time (21/30 trials) within 60 minutes.

##### Must Touch

This is similar to the Initial Touch stage, except that on incorrect response, no reward is delivered, and the trainer enters ITI. If no response is made in 2 minutes, the trial ends and enters ITI. This stage reinforces that only the stimulus is associated with the reward. Criterion is reached when 70% of the touches are correct (21/30 trials) within 60 minutes.

##### Must Initiate

This is similar to the Must Touch stage, except that the rodent is required to enter the reward port to ‘initiate’ the trial. This stage introduces initiation to minimize accidental touches. Criterion is reached when the rodent touches the correct stimulus 70% of the time (21/30 trials) within 60 minutes.

##### Punish Incorrect

This is similar to the Must Initiate stage, except if the incorrect response is selected, no reward is delivered and a 1s buzzer sounds. This stage strengthens discrimination learning using both positive reinforcement and punishment cues. Criterion is reached when the rodent touches the correct stimulus 70% of the time (21/30 trials) within 60 minutes over two consecutive days.

##### Simple Pairwise Discrimination

This is similar to the Punish Incorrect stage, except two new stimuli are presented (one correct, one incorrect). The trainer will enter correction trials (repeating the same trial) until the rodent picks the correct stimulus. This stage exposes the rodent to two new stimuli to learn and reduce any side bias through correction trials. Criterion is reached when the rodent touches the correct stimulus 70% of the time (21/30 trials) within 60 minutes over two consecutive days.

## 3. Results

### 3.1. Rat training

All rats successfully completed the full training pipeline advancing through all training stages. Learning performance in the last three stages: incorrect choice punishment, simple discrimination and novel discrimination are shown in Figure 7. All rats learned across the training timeline, with clear transitions between task phases as indicated by color-coded bars. As rodents improved in task performance, the number of correction trials—repeated presentations of the same stimulus pair following an incorrect choice—gradually decreased, reflecting improved discrimination accuracy.

**Figure 7.**
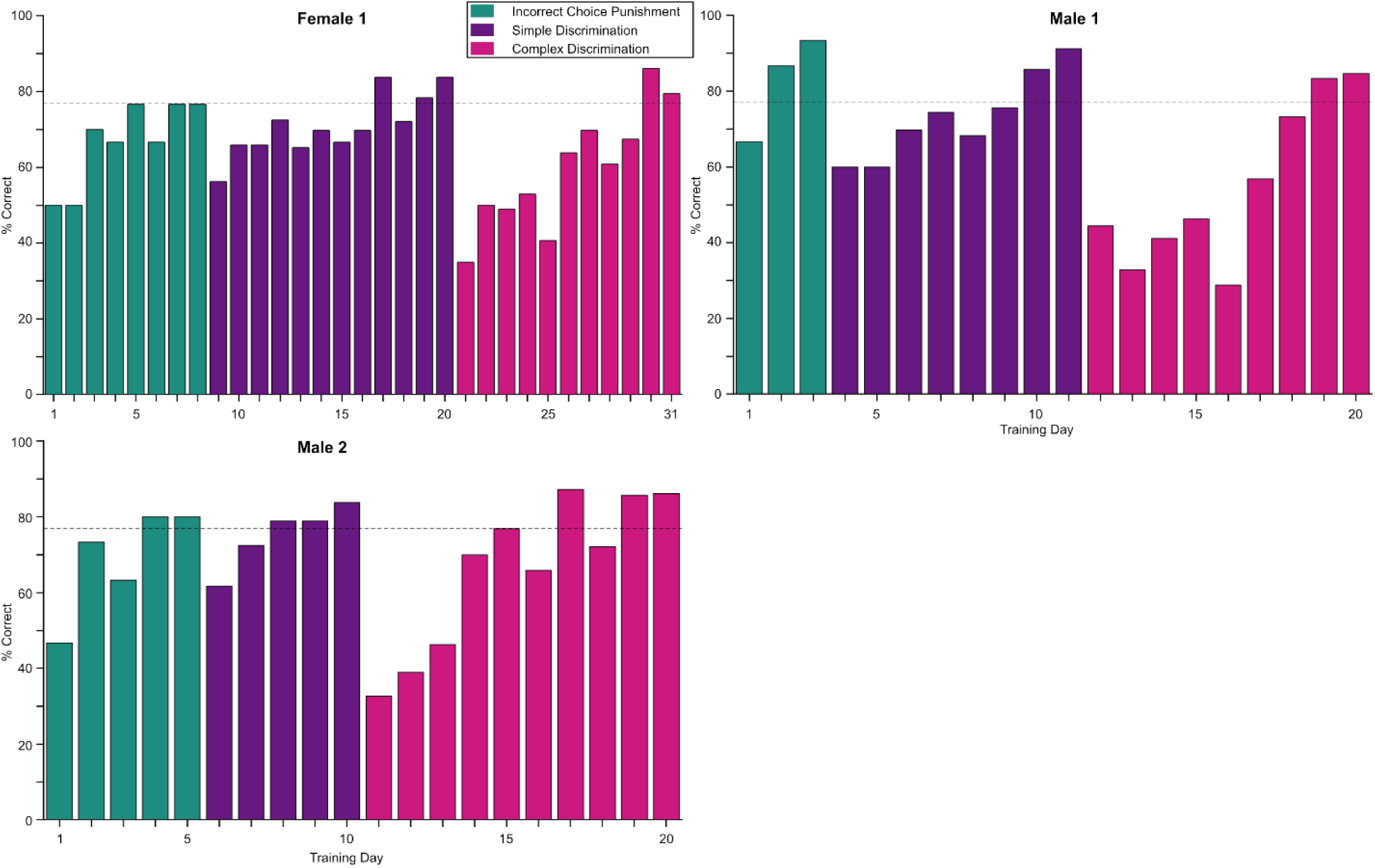
Rat learning curves. Daily percent-correct performance is shown for each rat (Male 1, Male 2, and Female 1) across the last three training stages. The dashed horizontal line marks the 77 percent accuracy criterion used to advance to the next stage.

### 3.2. Mouse Training

All 8 mice successfully completed the full training pipeline, advancing through all training stages. Most animals did not complete the “Must Initiate” stage because of time constraints but we chose to graduate them. Initiation was still required in all later stages and they were able to reach criterion quickly in the subsequent stage (Punish Incorrect). The mice reached criterion in the simple pairwise discrimination task in 35 ± 1.97 days (Fig. 8).

**Figure 8:**
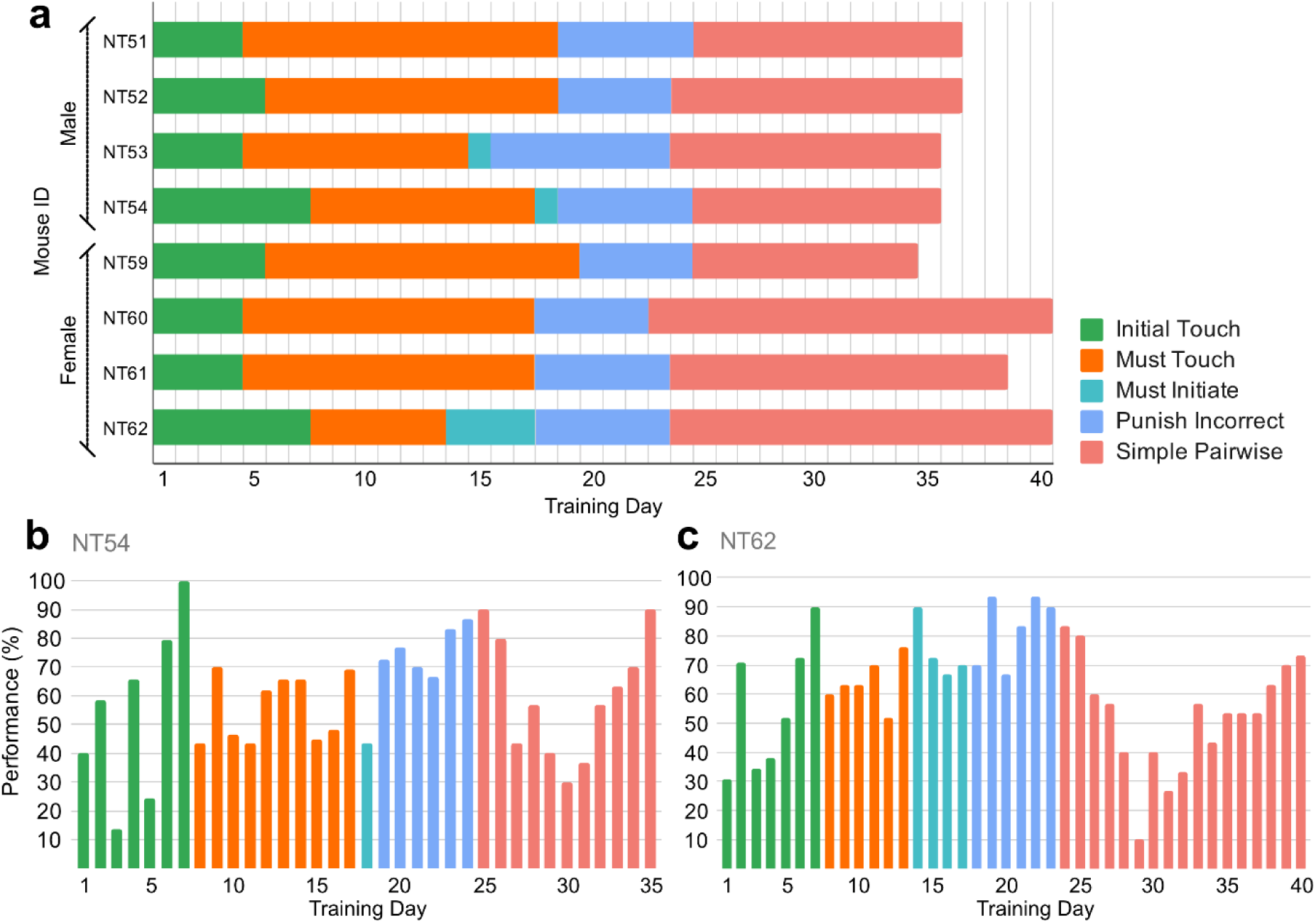
(a) Training timeline for the mice cohort, showing the number of days each mouse spent at each stage. Training stages are denoted by color coded bars. Data spans 40 days of training, after 8 days of habituation. **(b, c)** Performance of two example mice showing the success rate in each day of training.

### 3.3. Sound Attenuation

Figure 9 illustrates the results of our sound attenuation tests (for details, see Methods). Briefly, we ran tests across 6 different potentially sound-generating events as well as baseline, repeating each audio measurement (dB SPL) 5 times each under three different conditions: (1) Internal sound: measurement of events produced by a chamber within the same chamber, (2) Box Open: measurement of events produced by a chamber from a neighboring chamber, with the doors of sound attenuating boxes of both chambers open, and (3) Box Closed: same as Box Open, but with doors of both boxes open. Unsurprisingly, Buzzer and Reward pump actuation created noticeable auditory changes from baseline. In the box-closed condition, event sounds were suppressed to baseline levels (∼34dB across all conditions).

**Figure 9:**
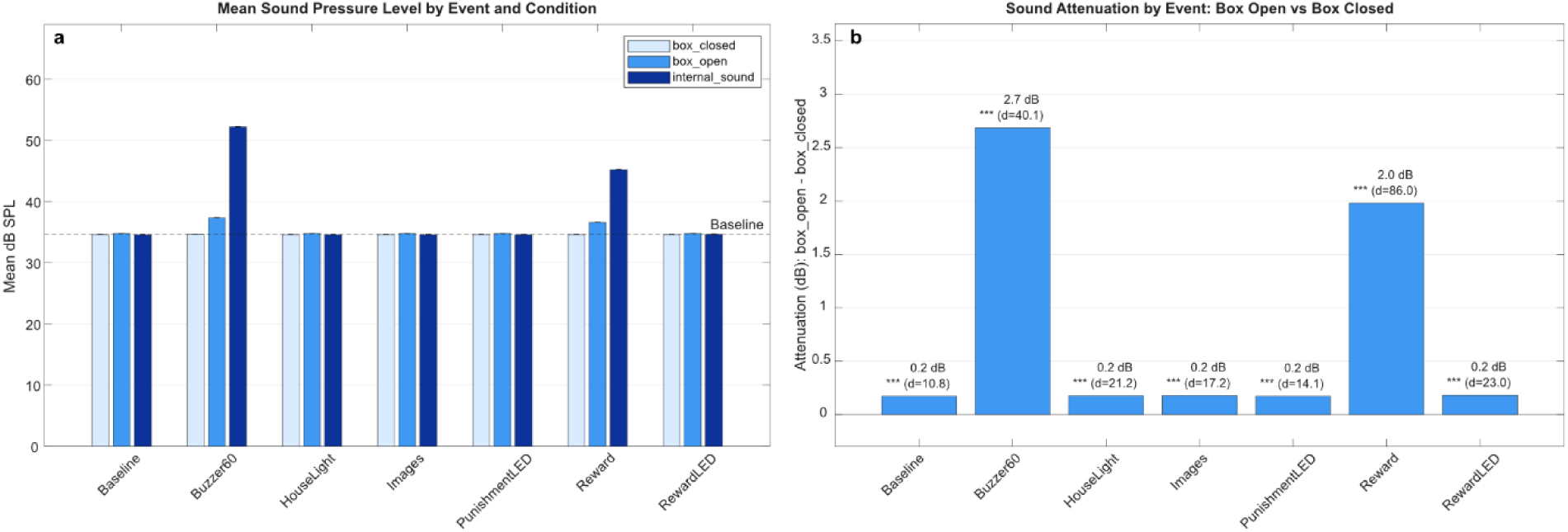
Sound attenuation results. **(a)** Sound pressure (Mean dB SPL) across Internal sound, Box Closed and Box Open conditions for 5 repeats of 7 events: operating the Buzzer, turning on House Light, turning on touchscreen Images, turning on Punishment LED, operating Reward Pump and turning on Reward LED. Error bars represent standard deviation across five trials and are very small due to the low variance. **(b)** Sound attenuation was quantified as the mean difference between box-open and box-closed conditions. Acoustic events showed significant attenuation, with the loudest event (Buzzer) showing the greatest attenuation (M = 2.69 dB, t4 = 89.68, p < .001, d = 40.11) and Reward showing moderate attenuation (M = 1.98 dB, t4 = 192.25, p < .001, d = 85.98). Presumed non-acoustic events (HouseLight, Images, PunishmentLED, RewardLED) showed negligible mean differences of 0.20 dB, though these also reached statistical significance (all t4 > 24.04, p < .001, d = 10.75–22.96).

## 4. Discussion

This work presents NC4Touch, an open-source, low-cost, modular and scalable touchscreen-based behavioral testing apparatus for mice and rats. Through successful training and validation in both species of rodents in a pairwise visual discrimination task, NC4Touch has proven to be a reliable platform for assessing learning and memory performance in a rodent model. The design’s modularity, affordability, and customizability make it a promising tool for laboratories seeking scalable and accessible solutions for cognitive testing in rodents.

A central feature of NC4Touch is its modular design. The use of independent touchscreen modules, a widely available controller (Raspberry Pi) and open-source software allows researchers to adapt the system to their own experimental needs. In addition, NC4Touch can be repaired, modified, and expanded by users. Either a single chamber or several chambers can be operated simultaneously, even running different, parallel tasks. The development of a virtual chamber simulation allows tasks to be tested and debugged before deploying on hardware. This flexibility is particularly important for laboratories that need to customize behavioral tasks, integrate additional sensors, or scale testing across multiple chambers while maintaining affordability.

We tested the NC4Touch system with both rats and mice, and found that male and female rodents of both species were able to learn a simple pairwise visual discrimination task and perform it with high accuracy. We also ran detailed sound attenuation tests that demonstrated that with chambers in isolation boxes lined with sound-absorbent sheets, auditory events from one chamber are indistinguishable from baseline sound level in neighboring chambers. Multiple chambers can thus be operated simultaneously in the same experimental room without risk of mutual behavioural interference.

Despite these strengths, several limitations should be acknowledged. First, while the apparatus accommodated mice and young adult rats, the current chamber dimensions may be restrictive for larger or older rats. However, because the CAD files are open-source and the system is modular, the chamber can be resized to fit different species, ages, or experimental requirements. Second, although the NC4Touch software framework is flexible and fully open-source, only a simple pairwise discrimination task has currently been developed and validated on the final version of the system. Additional behavioral paradigms will therefore need to be implemented and tested in future work. However, because the software is openly available and modular, users can design and customize their own behavioral tasks based on their experimental needs Third, like many DIY systems, NC4Touch requires some technical familiarity with electronics, software installation, and troubleshooting. While detailed documentation and assembly instructions are provided in our Open Science Framework page, sharing and integration with communities such as Touchscreen Cognition (https://touchscreencognition.org/) will be important for broader adoption.

Future directions for NC4Touch include the integration of automated pose estimation tools such as DeepLabCut^12^ or SLEAP^13^, which could allow for real-time behavioral tracking and more detailed analysis of movement patterns and response latencies. Additionally, incorporating automated head tracking and wireless neural recording / optogenetic platforms could expand the platform’s utility for systems neuroscience. Finally, using a single large capacitive touchscreen instead of multiple individual screens would simplify the design and increase flexibility.

Another important future direction is to broaden the range of validated cognitive tasks. At present, only visual pairwise discrimination has been tested using the NC4Touch. Future adaptations may include reversal learning, delayed match-to-sample, and object-location tasks, similar to those used in the Bussey-Saksida system^6^. Incorporating these paradigms would increase the platform’s utility for studying various domains of learning and memory.

NC4Touch also contributes to the growing open-source and DIY neuroscience movement, which aims to increase accessibility, reproducibility, and innovation through openly shared research tools and hardware. Similar community-driven initiatives, including Open Ephys for electrophysiology ^14^ and open-source miniature microscopes for in vivo imaging^15^, have demonstrated how low-cost and customizable technologies can accelerate neuroscience research while lowering financial barriers to entry. By providing openly available CAD files, parts lists, assembly instructions, and configurable software, NC4Touch aims to support similar collaborative development within the behavioral neuroscience community. Future software updates, hardware modifications, and additional behavioral paradigms will continue to be shared through online repositories and community resources.

Finally, NC4Touch offers a flexible, affordable, and open-source platform for rodent touchscreen-based behavioral testing. Its validation in both rats and mice, compatibility with multi-chamber operation, customizable software architecture, and virtual testing environment make it a valuable alternative to commercial touchscreen systems. By lowering financial and technical barriers, NC4Touch has the potential to support more accessible, reproducible, and collaborative behavioral neuroscience research.

## 5. Data and Code Availability

The design files for the NC4Touch chamber and a detailed assembly guide is available at our Open Science Framework page: https://osf.io/t5vfp

Software for operating the NC4Touch system, including the single chamber, multi-chamber and virtual chamber is available on our Github page: https://github.com/NC4Lab/NC4touch

## 6. Acknowledgments

This work was supported by a Kickstart Grant from the UBC Djawad Mowafaghian Centre for Brain Health (Snyder, Madhav), and two Natural Sciences and Engineering Research Council of Canada (NSERC) Discovery Grants (Snyder: RGPIN-2022-04468, Madhav: RGPIN-2024- 06614).

## 7. AI Disclosure

During the development of the NC4Touch software stack, the authors utilized GitHub Copilot, an AI-powered code completion tool, to aid in the generation, formatting, and refinement of custom scripts and state machine logic. All AI-assisted code was reviewed, iteratively tested, and rigorously validated by the authors to ensure accuracy.

## 8. Statement of Contributions

G.M. led the prototyping and development of the initial versions of the chamber, and developed the initial display and experimental control software. A.W.L. designed the hardware for the final chamber and supervised hardware assembly and iterations. M.B.C. developed the virtual chamber and contributed to the final version of chamber control and user interface code. I. S. performed initial rodent testing, devised the visual discrimination protocol with G.M., and assembled the OSF site and Bill of Materials. V.L. and J.Z. collected the mouse datasets, and generated results figures. L.C. collected the rat datasets. J. S. and M.S.M. supervised all trainees, guided the iterative design process, and edited the manuscript.

